# Analysis of Flurothyl-induced Seizures and Epileptogenesis in Mice with Targeted Deletions of Exons 3 and 4 in Dock7

**DOI:** 10.64898/2026.04.22.720243

**Authors:** Russell J Ferland, Talia Lizotte, Kathleen A Becker

## Abstract

Mutations in *DOCK7* have been identified in individuals with epileptic encephalopathies. Given that epileptic encephalopathies are a set of disorders that result in seizure activity and associated cognitive and behavioral impairments, we investigated the role of Dock7 in seizure susceptibility and flurothyl kindling using the repeated flurothyl seizure model in mice. Male and female *Dock7*^+/+^ and *Dock7*^△ex3-4/△ex3-4^ mice were subjected to 8 daily flurothyl exposures (kindling, induction phase) followed by a 28-day incubation period and a subsequent flurothyl rechallenge (retest). No significant differences were observed in baseline myoclonic jerk or generalized seizure thresholds between genotypes or sexes. However, over the kindling period, male *Dock7*^△ex3-4/△ex3-4^ mice exhibited slightly higher myoclonic jerk and generalized seizure thresholds compared to *Dock7*^+/+^ males across trials. Female mice showed similar trends, but the differences were only significant for generalized seizure thresholds. Following the 28-day incubation period and flurothyl retest, male mice of both genotypes maintained their seizure thresholds upon retest. *Dock7*^+/+^ female mice showed increased myoclonic jerk and generalized seizure thresholds during retest, while *Dock7*^△ex3-4/△ex3-4^ females maintained their thresholds. A key finding was the emergence of more severe forebrain→brainstem seizures upon flurothyl retest in a significant percentage of mice across all groups. However, the proportion of mice developing these seizures did not differ significantly between genotypes. Although *DOCK7* mutations have been linked to human epileptic encephalopathies and neurodevelopmental dysfunction, we find that *Dock7*^△ex3-4/△ex3-4^ male and female mice do not show heightened excitability or seizure susceptibilities using the repeated flurothyl seizure model.

**Highlights:**

- *Dock7*^△ex3-4/△ex3-4^ mice show slightly higher seizure thresholds during flurothyl kindling
- *Dock7*^△ex3-4/△ex3-4^ mice do not exhibit heightened seizure susceptibility upon retest.
- Forebrain-brainstem seizures emerged upon retest regardless of Dock7 genotype.

## 1. Introduction

Epilepsy affects approximately 1.2% of the U.S. population, making it one of the most prevalent neurological conditions (1). Current anti-epileptic treatments focus on suppressing seizures, but do not target the underlying mechanisms of epileptogenesis (2). While animal models have provided insights into seizure susceptibility, few studies have elucidated genes influencing epileptogenesis. Understanding the molecular pathways driving epileptogenesis represents an important research gap and an opportunity to develop new therapies. Recent genome-wide association studies (GWAS) in individuals with epilepsy have identified new genes associated with susceptibility to epilepsy (3-5). Moreover, genes for epilepsy have been identified through non-GWAS techniques, such as genetic mapping and whole exome sequencing (reviewed in (6, 7)). Specifically, *DOCK7* has been identified as a candidate gene for epilepsies, such as with epileptic encephalopathy (8-12). DOCK7 (dedicator of cytokinesis 7) is a member of the DOCK protein superfamily of guanine nucleotide exchange factors (GEFs) that activate small GTPases to regulate various cellular functions (13), particularly neural processes (14-17). Lastly, mice with a spontaneous mutation in *Dock7* have been previously shown to have no deficits in neurobehavioral function; however, measures of excitability or epilepsy were not assessed (18). Given the function of DOCK7 in human epilepsy, the mechanism by which DOCK7 dysfunction may contribute to epileptogenesis remains unclear. Here, we sought to determine whether mice with targeted deletions of exons 3 and 4 in *Dock7* (*Dock7*^△ex3-4/△ex3-4^) have differences in excitability and epileptogenesis, using the repeated flurothyl seizure model (19).

## 2. Materials and methods

### 2.1 Animals

*C57BL/6J-Dock7*^*em2/em2*^ (*Dock7*^△ex3-4/△ex3-4^) mice were generated as previously described (20, 21). These mice have global, homozygous deletions of exons 3 and 4 in *Dock7* and have abnormal trabecular bone acquisition and osteoblast function (21). For the seizure experiments described here, male and female *Dock7*^+/+^ and *Dock7*^△ex3-4/△ex3-4^ mice (n≥8 per group/sex) on a C57BL/6J background were produced through Dock7 matings in the University of New England (UNE) animal vivarium. Mice were maintained on a 12 h light-dark cycle with lights on at 7:00 AM with unlimited access to food and water. All protocols were performed under approval from the Institutional Animal Care and Use Committee of UNE and complied with the National Institutes of Health *Guide for the Care and Use of Laboratory Animals*.

When mice were at least 7–8 weeks of age, flurothyl seizure testing commenced. All seizure testing occurred between 8:00 AM and 12:00 PM. All animals were tested in the same chemical rated fume hood. The environment for seizure testing was maintained at a consistent temperature and humidity that did not differ significantly from day to day.

### 2.2 Flurothyl seizure induction (repeated flurothyl seizure model)

The repeated flurothyl seizure model (19) was used to assess seizure behaviors in mice [male *Dock7*^+/+^ (n = 9); male *Dock7*^△ex3-4/△ex3-4^ mice (n = 10); female *Dock7*^+/+^ (n = 11); female *Dock7*^△ex3-4/△ex3-4^ mice (n = 10)] (Fig.1). The total number of animals per group were run in three separate seizure cohort experiments. Mice were habituated to the experimental testing room for 30 minutes before each seizure trial. Then, each mouse was placed individually in a closed 2.4-liter Plexiglas chamber. A 10% flurothyl [Bis(2,2,2-trifluoroethyl) ether; Sigma Aldrich, St. Louis, MO] solution, made by diluting flurothyl in 95% ethanol, was infused into the chamber at a rate of 100μl/min onto a dry gauze pad suspended from the top of the chamber. Testing was conducted on one mouse at a time with a new gauze pad for each animal. As the flurothyl evaporated, mice were exposed to the vapors in the enclosed space. The first seizure phenotypes observed were myoclonic jerks, which are defined as brief, severe contractions of the neck and body musculature while maintaining postural control (22). Once mice lost postural control, indicating the start of a generalized seizure, the chamber was opened to room air. The generalized seizure threshold (GST) was defined as the latency from the start of flurothyl infusion to loss of postural control (23). Mice underwent one seizure trial per day for 8 days (induction phase). This was followed by a 28-day rest period without testing. On retest, a final flurothyl seizure was induced (rechallenge) to determine final seizure phenotypes. Multiple seizure phenotypes were quantified for each trial: 1) latency to the first myoclonic jerk (myoclonic jerk threshold), 2) number of myoclonic jerks preceding a generalized seizure, 3) latency to a generalized seizure (generalized seizure threshold), 4) duration of the generalized seizure, and 5) the behavioral seizure grade.

**Figure 1.**
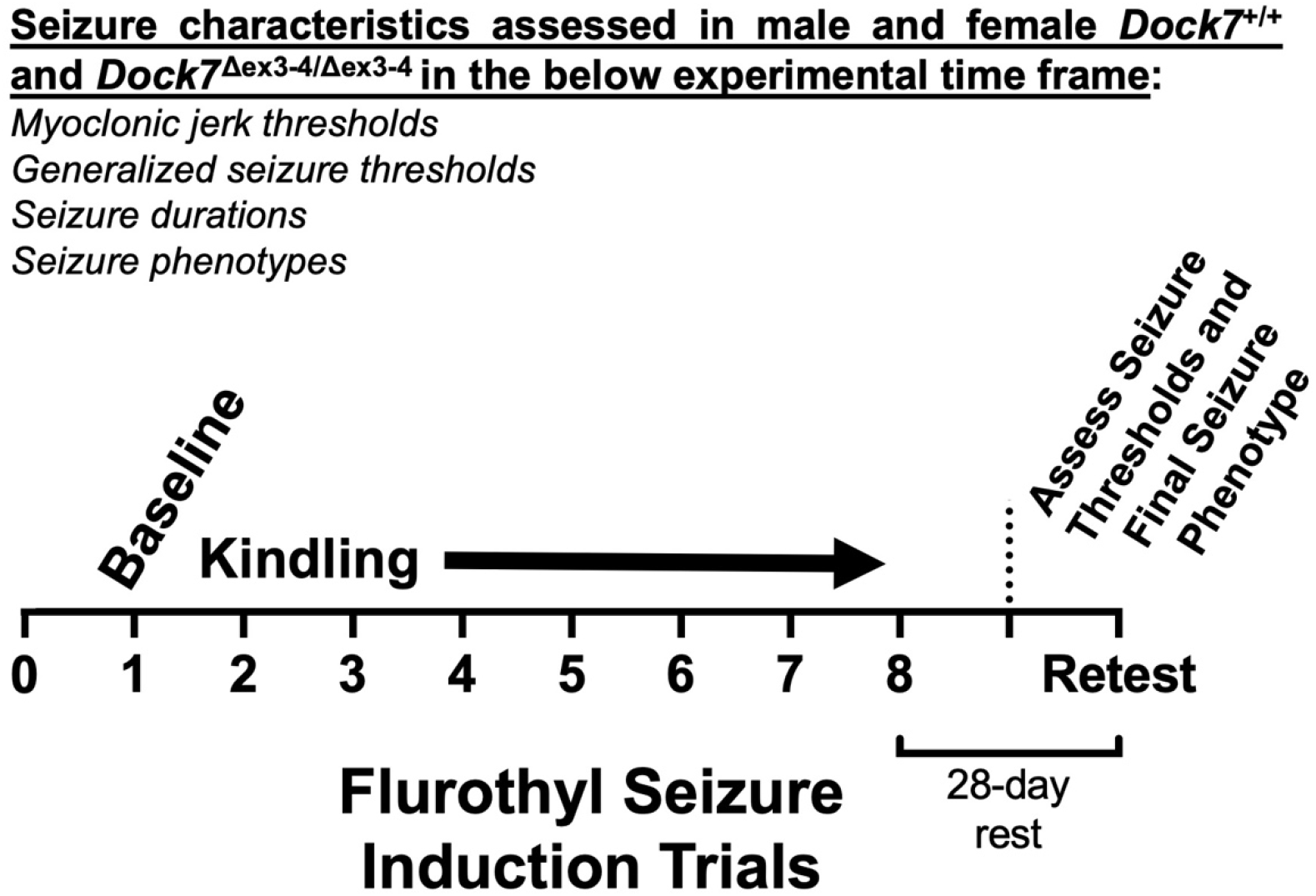
Diagram illustrating the experimental design. Mice are exposed to a 10% flurothyl solution, through evaporation-inhalation, in a closed chamber (19). In this model, the latency to the first myoclonic jerk as well as the latency to the initiation of a generalized seizure is determined in seconds and defines the myoclonic jerk threshold and generalized seizure threshold, respectively. The duration of each seizure is determined as well as the phonotype of the generalized seizure that is observed. The exposure to flurothyl occurs over 8-induction phase trials (kindling), followed by a 28-day retest trial. The first trial determines the baseline excitability for myoclonic jerk threshold and generalized seizure threshold. Repeated exposure to flurothyl over the 8 trials allows for determination of changes in seizure excitability, reminiscent of a “kindling” effect (decreases in seizure threshold [increases in excitability] with repeated seizures) (34). Importantly, the type of seizure expressed is classified, allowing for an understanding of potential seizure phenotype changes over these 8 trials. Following a 28-day rest period, mice are again exposed to flurothyl with the above seizure characteristics determined. This experimental model was utilized to assess whether there were any differences in these parameters in male and female *Dock7*^+/+^ and *Dock7*^△ex3-4/△ex3-4^ mice.

The behavioral seizure grading scale ranged from 1 to 7 based on seizure features indicative of forebrain versus brainstem ictal propagation, as previously described (19). Grades 1–2 are clonic-forebrain seizure behaviors, with loss of posture and facial/limb clonus, resembling electrically kindled generalized seizures (23). Grades 3–7 are tonic-brainstem seizures characterized by intense running, hopping, treading, and/or forelimb/hindlimb extension, reminiscent of tonic seizures seen in maximal corneal electroshock models (24). In summary, lower seizure grades (grades 1–2) reflect clonic seizures of forebrain origin, while higher grades (grades 3–7) reflect tonic seizures originating in brainstem regions. This standardized scale enabled tracking of seizure phenotypes over the induction and retesting protocol. Importantly, in the repeated flurothyl model, for seizures to reach grade 3 and above, the seizure must first start as a grade 1–2 seizure. As such, seizures induced in the forebrain seizure network can progress into the brainstem seizure network (a seizure referred to as a forebrain→brainstem seizure).

### 2.3 Statistical analysis

Within each sex, baseline (trial 1) measures of myoclonic jerk threshold, generalized seizure threshold, and seizure duration between *Dock7*^+/+^ and *Dock7*^△ex3-4/△ex3-4^ mice were analyzed using t-tests. Comparisons of myoclonic jerk thresholds, generalized seizure thresholds, and seizure durations over the 8 seizure trials utilized repeated measures ANOVAs, followed by Tukey Honest Significant Difference (HSD) post-hoc tests. Two-way repeated measures ANOVAs (with Tukey HSD post-hoc tests) were utilized to assess differences across seizure trials in comparing both genotype and sex. Paired t-tests compared trial 8 to the retest trial for each seizure characteristic. Fisher’s exact tests evaluated differences in the percentages of mice exhibiting forebrain (grades 1–2) versus forebrain→brainstem (grades 3–7) seizure types. All analyses were conducted using Statistica software (StatSoft, Tulsa, OK) or Prism (GraphPad, Boston, MA) with significance set at least *P* < 0.05.

## 3. Results

### 3.1 Effects of Dock7 on Baseline (Trial 1) Flurothyl-induced Myoclonic Jerk Thresholds

To investigate potential changes in baseline flurothyl-induced myoclonic jerk thresholds, male and female *Dock7*^+/+^ and *Dock7*^△ex3-4/△ex3-4^ mice were assessed. Analysis revealed no statistically significant differences in myoclonic jerk thresholds between genotypes for either sex (males: t_17_ = -0.45, *P* = 0.66; females: t_17_ = -1.31, *P* = 0.21). Additionally, no significant differences were found between sexes (F_3,36_ = 1.25, *P* = 0.31) (Fig. 2A).

**Figure 2.**
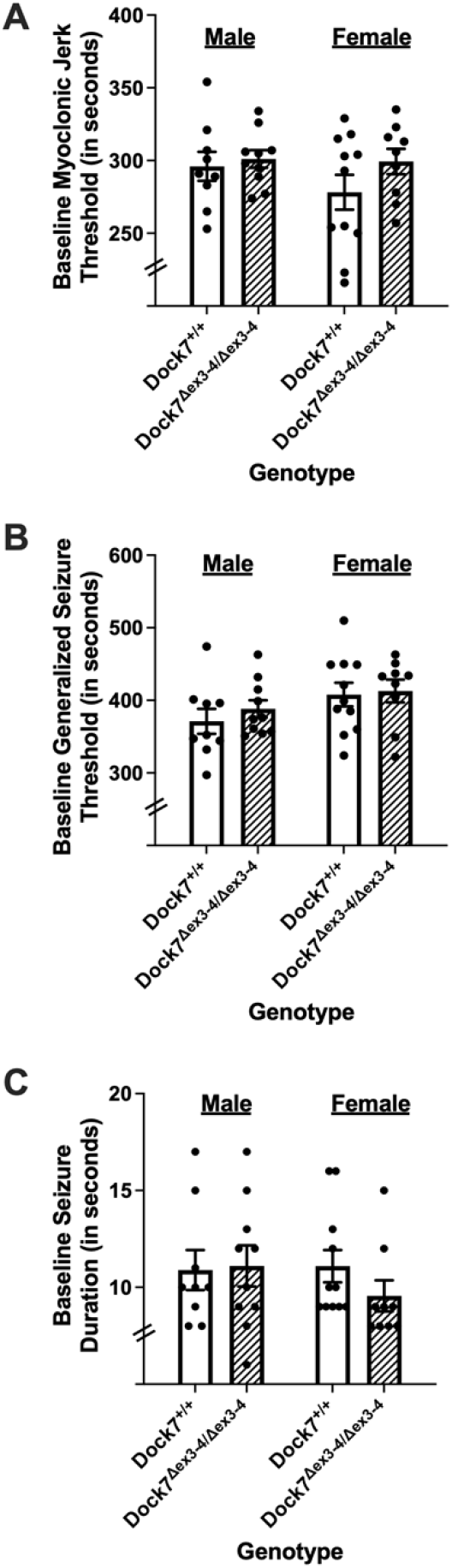
Baseline myoclonic jerk threshold, baseline generalized seizure threshold, and baseline seizure duration in *Dock7*^+/+^ and *Dock7*^△ex3-4/△ex3-4^ male and female mice. The latency (in seconds) to the first myoclonic jerk (myoclonic jerk threshold) (**A**), the latency (in seconds) to the first generalized seizure (generalized seizure threshold) (**B**), and seizure duration (in seconds) (**C**) was determined in male (blue) and female (pink) *Dock7*^+/+^ (open bars) and *Dock7*^△ex3-4/△ex3-4^ (cross-hatched bars) mice following exposure to 10% flurothyl. No statistically significant results were found between *Dock7*^+/+^ and *Dock7*^△ex3-4/△ex3-4^ mice for baseline myoclonic jerk threshold, baseline generalized seizure threshold, and baseline seizure duration. There were also no significant sex differences. Shown are the mean ± SEM for each group, with each mouse represented by an individual dot.

### 3.2 Effects of Dock7 on Baseline (Trial 1) Flurothyl-induced Generalized Seizure Thresholds and Seizure Durations

To determine whether *Dock7*^△ex3-4/△ex3-4^ mice had alterations in their flurothyl-induced generalized seizure thresholds or seizure durations, baseline flurothyl-induced generalized seizure thresholds (Fig. 2B) and seizure durations (Fig. 2C) were determined in male and female *Dock7*^+/+^ and *Dock7*^△ex3-4/△ex3-4^ mice. There were no statistically significant differences in generalized seizure thresholds between the *Dock7*^+/+^ and *Dock7*^△ex3-4/△ex3-4^ mice within each sex (males: t_17_ = -0.84, *P* = 0.41; females: t_18_ = -2.2, *P* = 0.83) or between sexes (F_3,35_ = 1.49, *P* = 0.23) (Fig. 2B). Similarly, no significant differences were found in seizure durations between the *Dock7*^+/+^ and *Dock7*^△ex3-4/△ex3-4^ mice within each sex (males: t_17_ = -0.14, *P* = 0.89; females: t_18_ = 1.31, *P* = 0.21) between sexes (F_3,35_ = 0.60, *P* = 0.62) (Fig. 2C).

### 3.3 Effects of Dock7 on Flurothyl-induced Myoclonic Jerk Thresholds over Eight Daily Seizure Trials (Kindling)

Repeated measures ANOVAs were used to determine whether there were differences in myoclonic jerk thresholds following 8 daily 10% flurothyl exposures in male and female *Dock7*^+/+^ and *Dock7*^△ex3-4/△ex3-4^ mice. Significant differences in myoclonic jerk thresholds were found between male *Dock7*^+/+^ and *Dock7*^△ex3-4/△ex3-4^ mice (F_1,17_ = 12.22, *P* = 0.003) with the male Dock7^+/+^ mice having slightly lower myoclonic jerk thresholds overall (Fig. 3A). In males, there were also the expected decreases in myoclonic jerk thresholds across the 8 flurothyl-induced seizure trials (F_7,119_ = 34.87, *P* < 0.0001) (Fig. 3A). There were no interaction effects with between genotype and seizure trials (*P* = 0.69) in male mice. In female mice, there were no significant differences in myoclonic jerk thresholds between *Dock7*^+/+^ and *Dock7*^△ex3-4/△ex3-4^ mice (F_1,19_ = 3.14, *P* = 0.093) (Fig. 3B). There were again the expected decreases in myoclonic jerk thresholds across the 8 flurothyl-induced seizure trials in female mice (F_7,133_ = 30.82, *P* < 0.0001) for both genotypes (Fig. 3B). There were no interaction effects with between genotype and seizure trials in female mice (*P* = 0.12). Lastly, when collapsing across both sexes and genotypes, there were no significant effects on myoclonic jerk thresholds in comparing genotype and sex.

**Figure 3.**
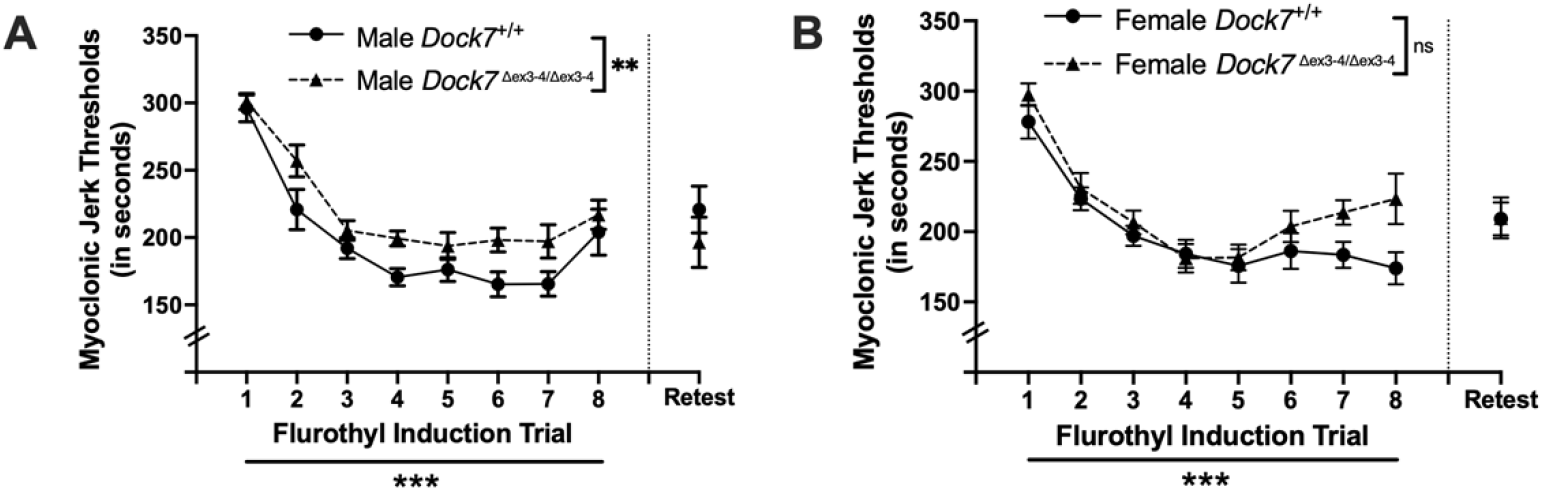
Changes in myoclonic jerk thresholds across eight seizure induction trials in *Dock7*^+/+^ and *Dock7*^△ex3-4/△ex3-4^ male and female mice. The latency (in seconds) to the first myoclonic jerk (myoclonic jerk threshold) after exposure to 10% flurothyl is shown for each flurothyl induction trial. Statistically significant decreases in myoclonic jerk thresholds across 8 flurothyl trials were observed in male (blue, **A**) and female (pink, **B**) *Dock7*^+/+^ and *Dock7*^△ex3-4/△ex3-4^ mice (***P* < 0.0001**). Significant differences were also observed between male *Dock7*^+/+^ and *Dock7*^△ex3-4/△ex3-4^ mice (***P* = 0.003**) for myoclonic jerk thresholds across the 8 seizure trials, with the male *Dock7*^△ex3-4/△ex3-4^ mice having slightly higher myoclonic jerk thresholds compared to the male *Dock7*^+/+^ mice (**A**). Conversely, there were no significant differences between female *Dock7*^+/+^ and *Dock7*^△ex3-4/△ex3-4^ mice for myoclonic jerk thresholds across the 8 seizure trials (**B**). Lastly, there were no significant interaction effects in myoclonic jerk thresholds between genotype and seizure trials for the male *Dock7*^+/+^ and *Dock7*^△ex3-4/△ex3-4^ mice and the female *Dock7*^+/+^ and *Dock7*^△ex3-4/△ex3-4^ mice, and no significant effects on myoclonic jerk thresholds when collapsing across sex and genotype. After a 28-day incubation period (indicated by a dashed vertical line), we conducted a flurothyl retest and compared myoclonic jerk thresholds to trial 8. In male mice, we observed no significant differences in myoclonic jerk thresholds for either *Dock7*^+/+^ or *Dock7*^△ex3-4/△ex3-4^ genotypes when comparing trial 8 to retest (**A**). Furthermore, there were no significant differences in myoclonic jerk thresholds between male *Dock7*^+/+^ and *Dock7*^△ex3-4/△ex3-4^ mice during the retest (**A**). For females, *Dock7*^△ex3-4/△ex3-4^ mice showed no significant differences in myoclonic jerk thresholds between trial 8 and retest (**B**). However, *Dock7*^+/+^ females exhibited a significant increase in myoclonic jerk threshold during the retest compared to trial 8. Despite this difference in *Dock7*^+/+^females, we found no significant differences in myoclonic jerk thresholds between female *Dock7*^+/+^ and *Dock7*^△ex3-4/△ex3-4^ mice during the flurothyl retest. Symbols denote the mean ± SEM for each group. \*\**P* = 0.003 (significance across the 8 trials between male *Dock7*^+/+^ and *Dock7*^△ex3-4/△ex3-4^ mice), \*\*\**P* < 0.0001 (significance across the 8 trials within genotype), ns=not significant

### 3.4 Effects of Dock7 on Flurothyl-induced Generalized Seizure Threshold over Eight Daily Seizure Trials (Kindling)

Repeated measures ANOVAs were used to assess differences in generalized seizure thresholds following 8 daily 10% flurothyl exposures in male and female *Dock7*^+/+^ and *Dock7*^△ex3-4/△ex3-4^ mice. In males, significant differences in generalized seizure thresholds were observed between *Dock7*^+/+^ and *Dock7*^△ex3-4/△ex3-4^ mice (F_1,17_ = 10.43, *P* = 0.005), with *Dock7*^*+/+*^ mice showing slightly lower generalized seizure thresholds overall (Fig. 4A). Both male genotypes exhibited the expected decrease in generalized seizure thresholds across the 8 flurothyl trials (F_7,119_ = 63.11, *P* < 0.0001) (Fig. 4A). No interaction effects between genotype and seizure trials were found in male mice (*P* = 0.64). Similarly, female mice showed significant differences in generalized seizure thresholds between genotypes, with *Dock7*^*+/+*^ females displaying slightly lower generalized seizure thresholds overall (F_1,19_ = 6.35, *P* = 0.021) (Fig. 4B). The expected decreases in generalized seizure thresholds across trials was also observed in females (F_7,133_ = 68.13, *P* < 0.0001) (Fig. 4B). No significant interaction effects between genotype and seizure trials were found in female mice (*P* = 0.82). Lastly, when data were combined across sex and genotype, no significant effects on generalized seizure thresholds were observed when comparing genotype and sex.

**Figure 4.**
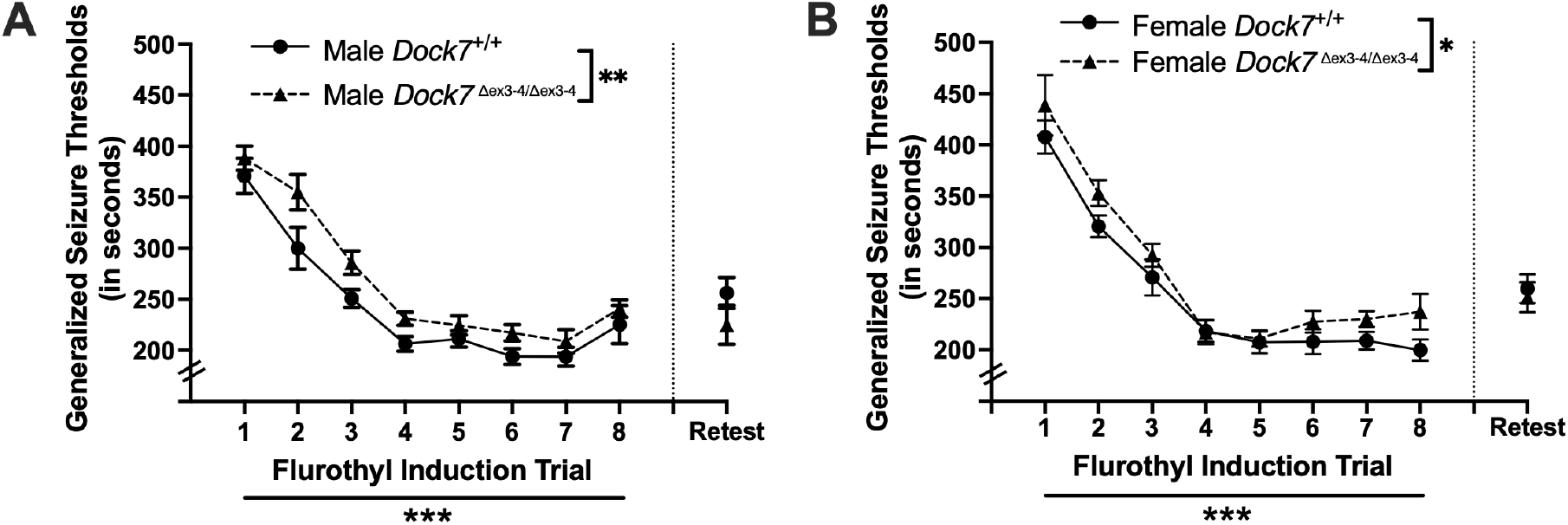
Changes in generalized seizure thresholds across eight seizure induction trials in *k7*^+/+^ and *Dock7*^△ex3-4/△ex3-4^ male and female mice. The latency (in seconds) to a generalized seizure (generalized seizure threshold) after exposure to 10% flurothyl is shown for each flurothyl induction trial. Generalized seizure thresholds decreased significantly across the 8 flurothyl trials in male (blue, **A**) and female (pink, **B**) mice in both genotypes, indicating kindling in all 4 groups (***P* < 0.0001**). Significant differences were also observed between male *Dock7*^+/+^ and *Dock7*^△ex3-4/△ex3-4^ mice ***P*= 0.005**) and female *Dock7*^+/+^ and *Dock7*^△ex3-4/△ex3-4^ mice (***P* = 0.021**) for generalized seizure thresholds across the 8 seizure trials, with both the male and female *Dock7*^△ex3-4/△ex3-4^ mice having slightly higher generalized seizure thresholds compared to their male and female *Dock7*^+/+^ mice, respectively (**A, B**). Lastly, there were no significant interaction effects in generalized seizure thresholds between genotype and seizure trials for the male *Dock7*^+/+^ and *Dock7*^△ex3-4/△ex3-4^ mice and the female *Dock7*^+/+^ and *Dock7*^△ex3-4/△ex3-4^ mice, and no significant effects on generalized seizure thresholds when collapsing across sex and genotype. Following a 28-day incubation period (indicated by a dashed vertical line), we conducted a flurothyl rechallenge (retest) and compared generalized seizure thresholds with trial 8 thresholds. In males, we observed no significant differences in generalized seizure thresholds for either *Dock7*^+/+^ or *Dock7*^△ex3-4/△ex3-4^ mice when comparing trial 8 to retest (**A**). Furthermore, there were no significant differences in generalized seizure thresholds between male *Dock7*^+/+^ and *Dock7*^△ex3-4/△ex3-4^ mice during the retest (**A**). For females, *Dock7*^△ex3-4/△ex3-4^ mice showed no significant differences in generalized seizure thresholds between trial 8 and retest (**B**). However, *Dock7*^+/+^ females exhibited a significant increase in generalized seizure threshold during the retest compared to trial 8. Despite this difference in *Dock7*^+/+^ females, we found no significant diffrences in generalized seizure thresholds between female *Dock7*^+/+^ and *Dock7*^△ex3-4/△ex3-4^ mice during the flurothyl retest (**B**). Symbols denote the mean ± SEM for each group. \**P* = 0.021, \*\**P* = 0.005, \*\*\**P* < 0.0001

### 3.5 Effects of Dock7 on Flurothyl-induced Seizure Durations over Eight Daily Seizure Trials

Repeated measures ANOVAs were utilized to assess differences in total seizure durations following 8 daily 10% flurothyl exposures in male and female *Dock7*^+/+^ and *Dock7*^△ex3-4/△ex3-4^ mice. For male mice, no significant differences in total seizure durations were observed between genotypes (F_1,17_ = 1.37, *P* = 0.258) (Fig. 5A). There were significant differences in seizure durations in males for both genotypes with repeated seizures, with seizure durations increasing with each trial (F_7,119_ = 9.60, *P* < 0.0001) (Fig. 5A). There were no interaction effects between genotype and seizure trials in male mice (*P* = 0.71). In female mice, significant differences in seizure durations were found between the *Dock7*^+/+^ and *Dock7*^△ex3-4/△ex3-4^ groups (F_1,18_ = 6.34, *P* = 0.022) (Fig. 5B). Seizure durations also significantly increased with repeated trials in female mice for both genotypes (F_7,126_ = 22.49, *P* < 0.0001) (Fig. 5B). Specifically, *Dock7*^△ex3-4/△ex3-4^ female mice initially had shorter seizure durations compared to *Dock7*^+/+^ mice on trials 1 and 2, but seizure durations increased from trials 3 to 8 (Fig. 5B). Accordingly, there was a significant interaction between genotype and seizure trial in female mice (F_7,126_ = 3.39, *P* = 0.002) (Fig. 5B). When combining data across sex and genotype, no significant effects on seizure duration were found when comparing genotype and sex.

**Figure 5.**
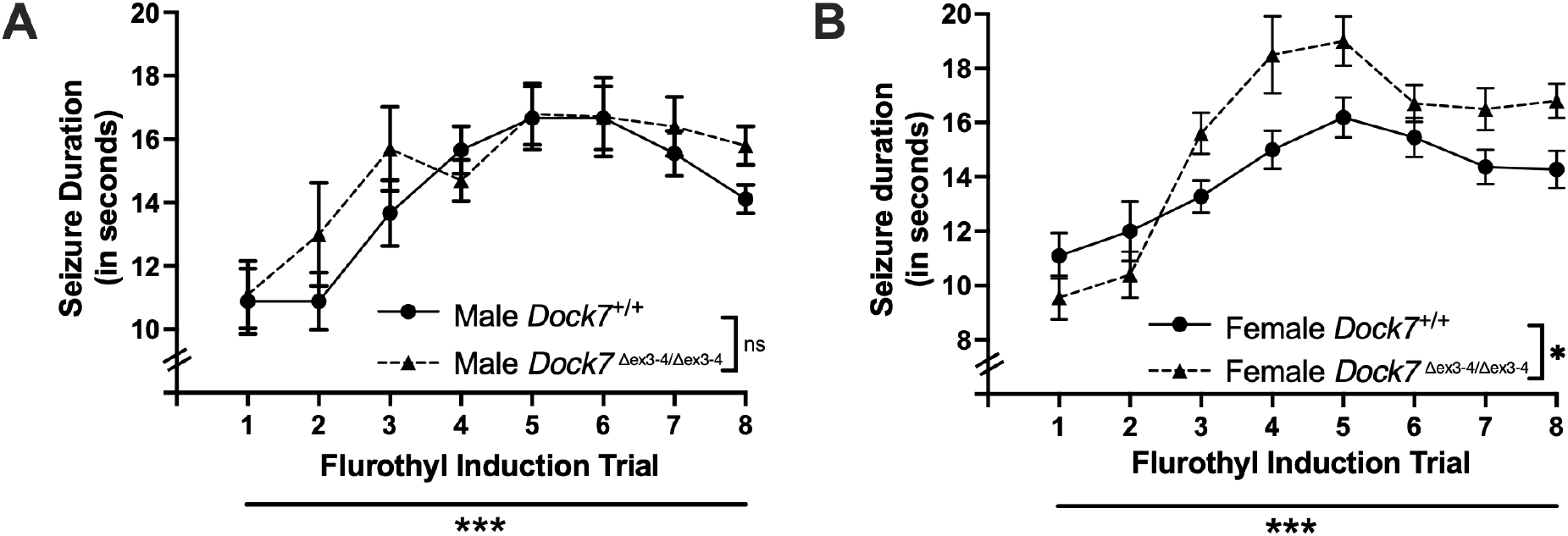
Changes in seizure durations across eight seizure induction trials in *Dock7*^+/+^ and *k7*^△ex3-4/△ex3-4^ male and female mice. The duration (in seconds) of each generalized seizure after exposure to 10% flurothyl is shown for each flurothyl induction trial. Statistically significant differences were found for both the male (blue, **A**) and female (pink, **B**) *Dock7*^+/+^ and *Dock7*^△ex3-4/△ex3-4^ mice in their seizure durations across the 8 flurothyl seizure induction trials (***P* < 0.0001**), demonstrating that all 4 groups of mice showed an increase in seizure durations with repeated seizures. In male mice, there were no significant differences between *Dock7*^+/+^ and *Dock7*^△ex3-4/△ex3-4^ animals in seizure durations across the 8 flurothyl seizure induction trials (**A**). However, there were significant differences in seizure duration between female *Dock7*^+/+^ and *Dock7*^△ex3-4/△ex3-4^ mice across the 8 trials (***P* = 0.022**), with female *k7*^△ex3-4/△ex3-4^ animals having longer seizure durations after trial 2 of the flurothyl induction phase (**B**). There were no significant interaction effects in seizure durations between genotype and seizure trials for the male *Dock7*^+/+^ and *Dock7*^△ex3-4/△ex3-4^ mice; however, there was a significant genotype by seizure trial interaction in female mice (***P* = 0.002**) (**B**). Lastly, no significant effects on seizure durations were found when collapsing across sex and genotype. Symbols denote the mean ± SEM for each group. \**P* = 0.022, \*\*\**P* < 0.0001

### 3.6 Effects of Dock7 on Comparisons of Trial Eight and Retest for Myoclonic Jerk Threshold, Generalized Seizure Threshold, and Seizure Duration

In this repeated flurothyl model, mice exposed to a retest of flurothyl following 8 flurothyl-induction trials and a 28-day incubation period have a maintenance of their myoclonic jerk and generalized seizure thresholds (22, 25). Comparisons within genotypes of trial 8 myoclonic jerk thresholds and trial 8 generalized seizure thresholds with retest myoclonic jerk thresholds and retest generalized seizure thresholds, respectively, revealed no significant differences in male myoclonic jerk (Fig. 3A) or generalized seizure (Fig. 4A) threshold. Additionally, there were no significant differences in myoclonic jerk thresholds (Fig. 3A) and generalized seizure thresholds (Fig. 4A) between *Dock7*^+/+^ and *Dock7*^△ex3-4/△ex3-4^ male mice on retest. In females, there were no differences in myoclonic jerk and generalized seizure thresholds in the *Dock7*^△ex3-4/△ex3-4^ mice in comparing trial 8 with retest (Fig. 3B and 4B). There were significant differences in both myoclonic jerk and generalized seizure thresholds in the *Dock7*^+/+^ mice comparing trial 8 and retest, with the retest myoclonic jerk (t_10_ = -2.48, *P* = 0.03) (Fig. 3B) and generalized seizure thresholds (t_10_ = -3.58, *P* = 0.005) (Fig. 4B) being higher than at the trial 8 timepoint. However, there were no significant differences in myoclonic jerk thresholds (Fig. 3B) and generalized seizure thresholds (Fig. 4B) between *Dock7*^+/+^ and *Dock7*^△ex3-4/△ex3-4^ female mice on flurothyl retest.

Mice tested in this repeated flurothyl model typically change their seizure phenotype following the incubation period and a flurothyl retest trial (25). This new seizure is longer in duration since it is typically composed of a clonic seizure that transitions into a seizure with tonic-brainstem manifestations. As such, both the male and female *Dock7*^+/+^ and *Dock7*^△ex3-4/△ex3-4^ mice had longer seizures on flurothyl rechallenge as compared to flurothyl induction trial 8 (*P* < 0.05; data not shown). No significant differences were noted in comparing seizure durations in *Dock7*^+/+^ and *Dock7*^△ex3-4/△ex3-4^ mice on flurothyl retest in both sexes.

### 3.7 Effects of Dock7 on the Alteration in Behavioral Seizure Phenotype in Response to Flurothyl-Induction Trials and Flurothyl Retest

All male and female *Dock7*^+/+^ and *Dock7*^△ex3-4/△ex3-4^ mice expressed clonic-forebrain seizures during the 8-day flurothyl induction trials, with no mice having forebrain→brainstem seizures (clonic seizures that transition into seizures with tonic (brainstem) manifestations) (Fig. 6). However, following this 8-day flurothyl induction phase and a 28-day incubation phase (during which no flurothyl-induced seizures were elicited), a significant percentage of male and female *Dock7*^+/+^ and *Dock7*^△ex3-4/△ex3-4^ mice expressed the expected new phenotype of a forebrain→brainstem seizure upon flurothyl rechallenge.

**Figure 6.**
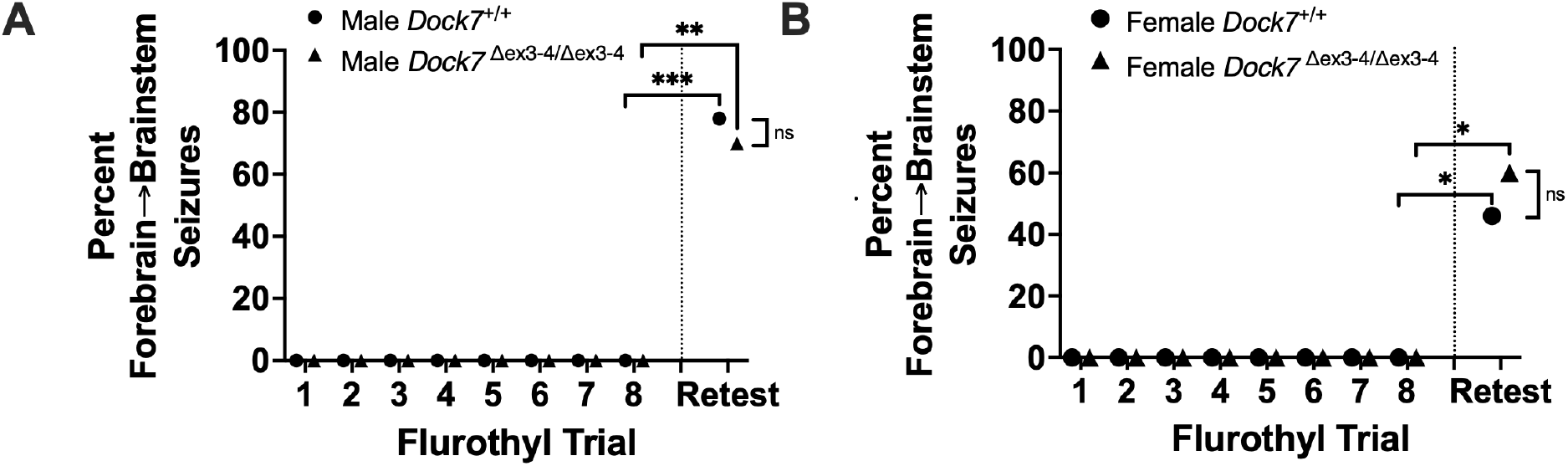
Behavioral seizure phenotypes across eight flurothyl seizure induction trials, and a 28-day incubation and flurothyl rechallenge, in *Dock7*^+/+^ and *Dock7*^△ex3-4/△ex3-4^ male and female mice. During the flurothyl-induction phase (trials 1-8), all male (blue, **A**) and female (pink, **B**) *Dock7*^+/+^ and *Dock7*^△ex3-4/△ex3-4^ mice express clonic, forebrain-mediated seizures upon inhalant exposure to flurothyl, with no forebrain→brainstem seizures being observed. However, following a 28-day incubation phase (denoted by the dashed vertical line) and flurothyl rechallenge (retest), we observed a marked increase in forebrain-brainstem seizures across both genotypes and sexes, with mice now exhibiting clonic (forebrain) seizures that transition uninterruptedly into seizures with tonic (brainstem) manifestations. Fisher’s Exact tests were used to determine that 78% of *Dock7*^+/+^ and 70% of *Dock7*^△ex3-4/△ex3-4^ male mice exhibited this seizure type, with both groups showing significant increases (***P* = 0.0007** and ***P* = 0.0031**, respectively) (**A**). Similarly, female mice demonstrated an increased incidence of forebrain→brainstem seizures, with 46% of *Dock7*^+/+^ and 60% of *Dock7*^△ex3-4/△ex3-4^ mice affected (***P* = 0.035** and ***P* = 0.011** respectively) (**B**). Notably, we found no significant differences in the percentage of forebrain→brainstem seizures between *Dock7*^+/+^ and *Dock7*^△ex3-4/△ex3-4^ mice within either sex group during the retest (**A, B**). These results suggest that while repeated flurothyl exposure increases susceptibility to forebrain→brainstem seizures, the Dock7 dysfunction does not significantly influence this susceptibility. Symbols denote the percentage of mice expressing forebrain→brainstem seizures for each group. \**P* < 0.04, \*\**P* < 0.004, \*\*\**P* < 0.0008, ns = not significant

Upon flurothyl retest, 78% of male *Dock7*^+/+^ mice and 70% of male *Dock7*^△ex3-4/△ex3-4^ mice expressed a forebrain→brainstem seizure (*Dock7*^+/+^; *P* = 0.0007 and *Dock7*^△ex3-4/△ex3-4^; *P* = 0.0031) by Fisher’s Exact test (Fig. 6A), with no significant differences observed between genotypes (Fig. 6A). In female mice on flurothyl retest, 46% of female *Dock7*^+/+^ mice and 60% of female *Dock7*^△ex3-4/△ex3-4^ mice expressed a forebrain→brainstem seizure (*Dock7*^+/+^; *P* = 0.035 and *Dock7*^△ex3-4/△ex3-4^; *P* = 0.011) by Fisher’s Exact tests (Fig. 6B). There were no significant differences in the percentage of forebrain→brainstem seizures between genotypes in female mice on flurothyl retest (Fig. 6B).

These findings indicate that repeated flurothyl exposure enhances the likelihood of forebrain→brainstem seizures in both *Dock7*^*+/+*^ and *Dock7*^△ex3-4/△ex3-4^ mice. However, the absence of significant differences between genotypes suggests that deletion of exons 3 and 4 in *Dock7* does not play a crucial role in modulating this increased susceptibility to more severe seizure types following exposure to the repeated flurothyl model.

## 4. Discussion

Epileptic encephalopathies are severe neurological disorders characterized by frequent seizures that significantly impair cognitive and behavioral development (26). Given that mutations in *DOCK7* have been identified in individuals with epileptic encephalopathies (8-10, 12, 27, 28), we hypothesized that *Dock7*^△ex3-4/△ex3-4^ mice would exhibit increased seizure susceptibility and more severe seizure phenotypes using the repeated flurothyl seizure model. Unexpectedly, *Dock7*^△ex3-4/△ex3-4^ mice did not show alterations in baseline flurothyl-induced seizure characteristics. Contrary to further expectations, *Dock7*^△ex3-4/△ex3-4^ mice showed slightly higher seizure thresholds during flurothyl kindling than *Dock7*^*+/+*^ mice, with males more affected than females. After a 28-day incubation, both genotypes developed more severe seizures upon retest, with no significant differences found between the groups. Despite the association of DOCK7 with human epileptic encephalopathies, our findings suggest that *Dock7*^△ex3-4/△ex3-4^ mice do not exhibit increased excitability or seizure susceptibility in this model, highlighting the complex relationship between gene mutations and epilepsy phenotypes.

The fact that *Dock7*^△ex3-4/△ex3-4^ mice did not have baseline differences from *Dock7*^+/+^ mice in myoclonic jerk thresholds, generalized seizure thresholds or seizure durations was surprising, given that *DOCK7* mutations give rise to epileptic encephalopathies in humans. Moreover, the repeated flurothyl model used here has been very sensitive in dissecting differences in seizure susceptibilities, even between different inbred strains of mice (29-31). The lack of significant differences in these baseline seizure characteristics between *Dock7*^*+/+*^ and *Dock7*^△ex3-4/△ex3-4^ mice indicates that *Dock7* dysfunction in mice does not inherently predispose them to increased seizure susceptibility.

Also unexpected were the marginally higher myoclonic jerk and generalized seizure thresholds observed in male *Dock7*^△ex3-4/△ex3-4^ mice, and the slightly higher generalized seizure thresholds observed in female *Dock7*^△ex3-4/△ex3-4^ mice, during the kindling (trials 1–8) (23). These results would indicate a potential protective effect of *Dock7* dysfunction with flurothyl kindling. However, it is important to note that while these differences were statistically significant, this observed protective effect was minimal.

The emergence of more severe forebrain→brainstem seizures upon flurothyl-induction, incubation (28 days), and flurothyl retest across all groups, regardless of genotype, suggests that this progression is likely independent of Dock7 function. This observation underscores the multifaceted nature of epileptogenesis and the likely involvement of multiple genetic and environmental factors. Moreover, some of the sex-specific differences observed in this model are interesting and warrants further investigation into the interaction between Dock7 and sex hormones or sex-specific neural circuits.

The discrepancy between our findings in mice and the association of *DOCK7* mutations with human epileptic encephalopathies raises several important points for consideration. The *Dock7*^△ex3-4/△ex3-4^ mice (21, 32) used in this study were generated from a *Dock7* floxed model that was developed using CRISPR/Cas9 to flank exons 3 and 4 of *Dock7* with LoxP sites (20). Deletion of exons 3-4 in *Dock7* is predicted to produce a premature stop codon; however, molecular analysis revealed the continued presence of the *Dock7* transcript and Dock7 protein, even though exons 3–4 in *Dock7* were eliminated at the transcript level (21). Our *Dock7* conditional mice, with deletion of exons 3-4, have been used in other deletion studies and have demonstrated a reduction in Rac1 activation with deletion of *Dock7* exons 3-4 (32). As such, *Dock7*^△ex3-4/△ex3-4^ mice may retain partial Dock7 function rather than representing a true null mutant. Further studies are necessary to assess changes in Dock7 function. Importantly, *Dock7*^△ex3-4/△ex3-4^ (homozygous) mice exhibited a diluted coat color, white belly spot, and abnormal bone phenotypes consistent with spontaneous Dock7 mutant mice (18, 20, 33). These phenotypes were not present in *Dock7*^△ex3-4/+^ (global heterozygous) mice (21). The absence of a robust seizure phenotype in *Dock7*^△ex3-4/△ex3-4^ mice may reflect this partial disruption of Dock7 function. However, our results clearly demonstrate that exons 3–4 of *Dock7* are not critical for seizure susceptibility or epileptogenesis, at least in the flurothyl seizure model.

The discrepancy between our findings and the association of *DOCK7* mutations with human epileptic encephalopathies may also reflect species-specific differences in how Dock7 influences neuronal excitability and seizure susceptibility. Moreover, it is also possible that the expected seizure differences could have been observed if different aged mice were examined, as in the current study mice were 7–8 weeks old at the beginning of the seizure testing.

## Acknowledgments

This work was partly supported by NIH grants R16NS140309 (to RJF) and P20GM152330 (to KAB).

## CRediT authorship contribution statement

**Russell J. Ferland**: Conceptualization, Formal analysis, Investigation, Resources, Data curation, Writing – original draft, Visualization, Supervision, Project administration, Funding acquisition. **Talia Lizotte**: Investigation, Resources, Writing – review & editing. **Kathleen A Becker**: Conceptualization, Resources, Writing – review & editing, Supervision, Project administration, Funding acquisition.

